# Pharmacological Evaluations of Select Herbal Extracts on TLR7/8-induced Cytokine and Chemokine Production in Macrophage-like Cells

**DOI:** 10.1101/2023.07.16.549251

**Authors:** Manisha Dagar, Kamala Priya, Madhu Dikshit, Ajay Kumar

## Abstract

Inflammation is an innate immune response triggered by harmful stimuli, such as pathogens, tissue injury, or toxins. The purpose is to eliminate the source of infection and initiate the healing process. However, an excessive acute inflammatory response can lead to severe and life-threatening complications, as seen during recent pandemics. In the context of viral infections, the activation of the TLR7/8 signaling pathway has been implicated in excessive cytokine secretion. In this study, we aimed to replicate the perturbed inflammatory environment resulting from the activation of the TLR7/8 specific agonists, imiquimod, and resiquimod.

We utilized macrophage-like cells, as macrophages are the first responders during infections and secrete key pro-inflammatory cytokines (TNF-α, IL-6, and IL-1β) to recruit immune cells to the site of infection. Herbal medicines have been traditionally used for centuries to enhance respiratory immune function. In the present study, we employed a prophylactic approach, where macrophage-like THP1 cells, differentiated with PMA, were pre-treated with select herbal extracts/formulations prior to TLR7/8 activation in the presence of agonists.

Several medicinal plants and formulations known for their therapeutic potential in respiratory ailments were investigated, including *Withania somnifera, Tinospora cordifolia, Glycyrrhiza glabra,* and AYUSH-64, an herbal formulation. The gene expression and corresponding secreted levels of various inflammatory mediators were measured using RT-PCR and ELISA methods, respectively, in treated and untreated differentiated THP1 cells induced with TLR7/8 agonists. Comparatively, the gene expression of inflammatory markers was significantly higher in resiquimod-induced cells than in imiquimod-treated cells. Notably, *Withania somnifera* demonstrated pronounced prophylactic efficacy compared to other herbs/formulations, as evidenced by reduction in expression of majority of investigated inflammatory marker genes.

## INTRODUCTION

Excessive, acute, and uncontrolled release of pro-inflammatory cytokines into circulation during the recent COVID-19 pandemic resulted in severe life-threatening complications and even death in many cases. Such a storm of cytokines may lead to an increased flux of immune cells (macrophages, neutrophils, and T cells) to the site of infection eventually causing more damage than repair [30]. Herbal therapies have been used in traditional medicine for centuries and are also being explored for their potential benefits in cytokine storm management. While the research on specific herbal treatments for cytokine storm is limited, some herbs have shown immunomodulatory and anti-inflammatory properties that may help to alleviate the excessive immune response.

Administration of TLR7/8 agonists, such as imiquimod or resiquimod, has been associated with the development of cytokine storm [19,9]. Among others, TNF-α, IL-6, and IL-1β are the key pro-inflammatory cytokines which drive the excessive secretion of cytokines. Their increased levels are often correlated with the severity of cytokine storm which may eventually lead to tissue damage and organ dysfunction [13,36,20]. TNF-α has been correlated to pro-inflammatory response mediated by IL-1β and IL-6. It is found to be involved in the regulation of inflammatory responses, infectious diseases, and malignant tumors [24]. Higher levels of TNF-α were found in severe COVID-19 patients [13]. Among other immune cells, monocytes and macrophages are the main source of these pro-inflammatory cytokines [7]. Human THP1 cells are monocyte-derived macrophage-like cells which mimic the macrophage functions and therefore are routinely used in *in-vitro cell* models in basic research. Such cell-based models were instrumental during the recent pandemic where they not only helped in disease understanding but were also the first line preclinical models to screen a large number of drug candidates for their efficacy and safety. THP1 cells differentiate into macrophage-like cells when incubated in the presence of PMA. TLR7/8 specific signalling was induced in the presence of specific agonists and measured in terms of gene expression as well as circulating levels of various inflammatory genes.

We have evaluated a few select herbal extracts that have traditionally been used over centuries in therapeutics to cure different infectious as well as immunocompromised conditions. A similar line of investigations has been reported with other herbs [2] or herbal oils [26] using different disease models. Mukherjee et al submitted that *Syzygium aromaticum, Piper nigrum, Cimicifuga racemosa, Zanthoxylum asiaticum, Camellia sinensis, Curcuma longa,* and *Tribulus terrestris* were effective in Covid-19 therapeutics. Similarly, the use of oil formulations such as Anu oil and Til tailya in the management of SARS-CoV-2 was explored by Rizvi et al [26] where the authors observed a significant reduction in viral load as an effect of treatment with Anu oil.

In the present investigation, we have specifically examined the efficacy of *Withania somnifera, Tinospora cordifolia, Glycyrrhiza glabra*, and AYUSH-64 in mitigating the inflammatory response triggered by TLR-7/8 induction in the presence of the ligands imiquimod and resiquimod. While the first three are individual herbs, AYUSH-64 is an herbal formulation comprising a combination of four different herbs (*Alstonia scholaris*, *Caesalpinia crista*, *Picrorhiza kurroa* and *Swertia chirata*), each contributing to the overall therapeutic benefits. To assess the prophylactic effects of these herbs/formulations, we employed a macrophage-like cell line model. Our objective was to identify the herb or formulation that exhibited the highest efficacy in either suppressing the TLR7/8-induced inflammatory response by promoting an anti-inflammatory environment. We evaluated various inflammatory gene markers, including cytokines and chemokines, both at the gene expression level and their secreted levels in the supernatant. To analyse gene expression, we extracted RNA from the cell lysates and employed the RT-PCR method. Simultaneously, we measured the secreted cytokine levels in the culture supernatant using the ELISA method. These analyses enabled us to determine the impact of the tested herbs/formulations on the expression of inflammatory genes and the corresponding secretion of cytokines.

## METHODOLOGY

### Cell culture

THP1 is a spontaneously immortalized monocyte-like cell line, derived from the peripheral blood of a child suffering from acute monocytic leukemia [33]. Culture conditions for the THP1 cell line were as described by ATCC. Briefly, RPMI 1640 (Gibco) supplemented with 10% FCS (Hyclone) and 1x penicillin (stock concentration = 100U/mL), and streptomycin (stock concentration = 100μg/mL) were used to culture the cell line. Cells were maintained in complete media at 37°C in humidified conditions with 5% CO_2_. Trypan blue was used to confirm cell viability. THP1 cells were seeded in the presence of PMA (5ng/mL) and allowed to differentiate for 48 hours and subsequently used.

### Preparation of the herbal extracts/formulations

Powdered form of well-characterized herbal extracts - WS, TC, GG and A64, prepared under the GMP conditions [17], were obtained from the National Medicinal Plants Board (NMPB), Ministry of AYUSH, Government of India. A stock solution of 4mg/mL was prepared in the RPMI medium. RPMI medium (1mL) was added to 4mg of each extract/formulation powder and kept in dark for overnight shaking at room temperature. Subsequently, the extract solutions were centrifuged at 10,000 rpm for 10 minutes and the supernatant was passed through a 0.22μm syringe filter in sterile environment. This supernatant was assumed to be an absolute concentrate of the herbal extract and was used for investigations in this study. All stocks were prepared fresh for each experiment and the filtered stocks were saved at 4°C in dark.

### Cell Viability Assay

Cell viability in the presence and absence of herbal extracts/formulations was evaluated by MTT assay. Differentiated THP1 cells were treated with various concentrations of herbal extracts (100, 300, and 1000μg/mL). After 24 hours of treatment, cells were incubated in dark with 0.5mg/mL MTT (Sigma, Cat no # M2128) for 4 hours at 37°C. The culture supernatant was gently removed and replaced with 100μL DMSO per well (Sigma, Cat no # 276855). The cells were allowed to incubate in dark at RT with constant shaking for 30 minutes. Subsequently, the absorbance was measured with a microplate reader (Molecular Devices, Spectramax M2) at 570nm wavelength. Dexamethasone (Selleck Chemicals, # S1322) (1μM) was used as an internal reference. Latter is a steroidal drug with powerful anti-inflammatory properties.

### qPCR to study gene expression of Inflammatory mediators

Differentiated THP1 cells (1 million cells/well), seeded in a 6-well plate, were pre-treated for 60 minutes with herbal extracts/formulations at different concentrations (3μg/mL, 10μg/mL, 30μg/mL, 100μg/mL, and 300μg/mL). Subsequently, TLR7/8 agonists (Imiquimod (Tocris # 3700/50) / Resiquimod (Sigma # SML1096), respectively) were added to induce TLR signalling. Post induction, the cells, and culture supernatants were collected separately. Total RNA was extracted using a Trizol reagent (Sigma # T9424). Qualitative and quantitative RNA analysis was done on a Multiskan GO plate reader (Thermo Scientific). Reverse transcription of 250ng RNA was done using the high-capacity cDNA reverse transcription kit (Applied Biosystems # 4368814). For the real-time quantitative PCR, the cDNA was amplified using PowerUp SYBR Green mix (Applied Biosystems # A25742) following the standard protocol for the dye on QuantStudio 6 Real-Time PCR System (Applied Biosystems). Cycling conditions were UDG activation at 50°C for 2 minutes, Hold Dual-Lock™ DNA polymerase at 95°C for 2 minutes, and 40 cycles of Denaturation at 95°C for 15 seconds and annealing/extension at 60°C for 1 minute each. For each reaction 300nM of forward and reverse primer mix was used (Table 1). The relative mRNA expression data were analyzed using the ΔΔCt method and the values were normalized to beta-actin mRNA levels. The data was expressed with respect to the fold change as compared to the untreated control.

**Table 1:**
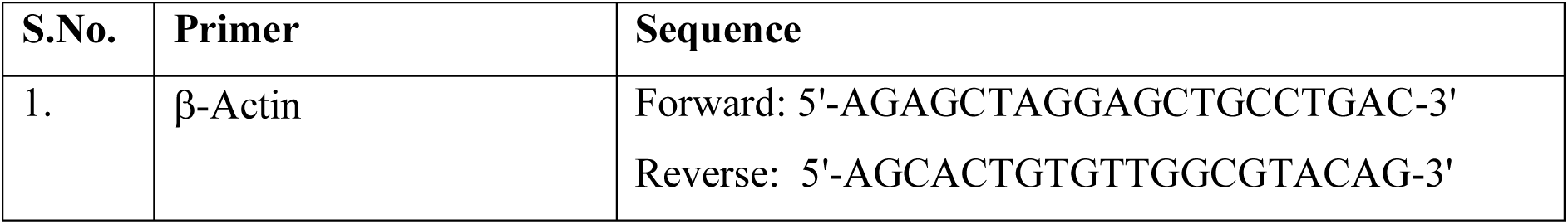
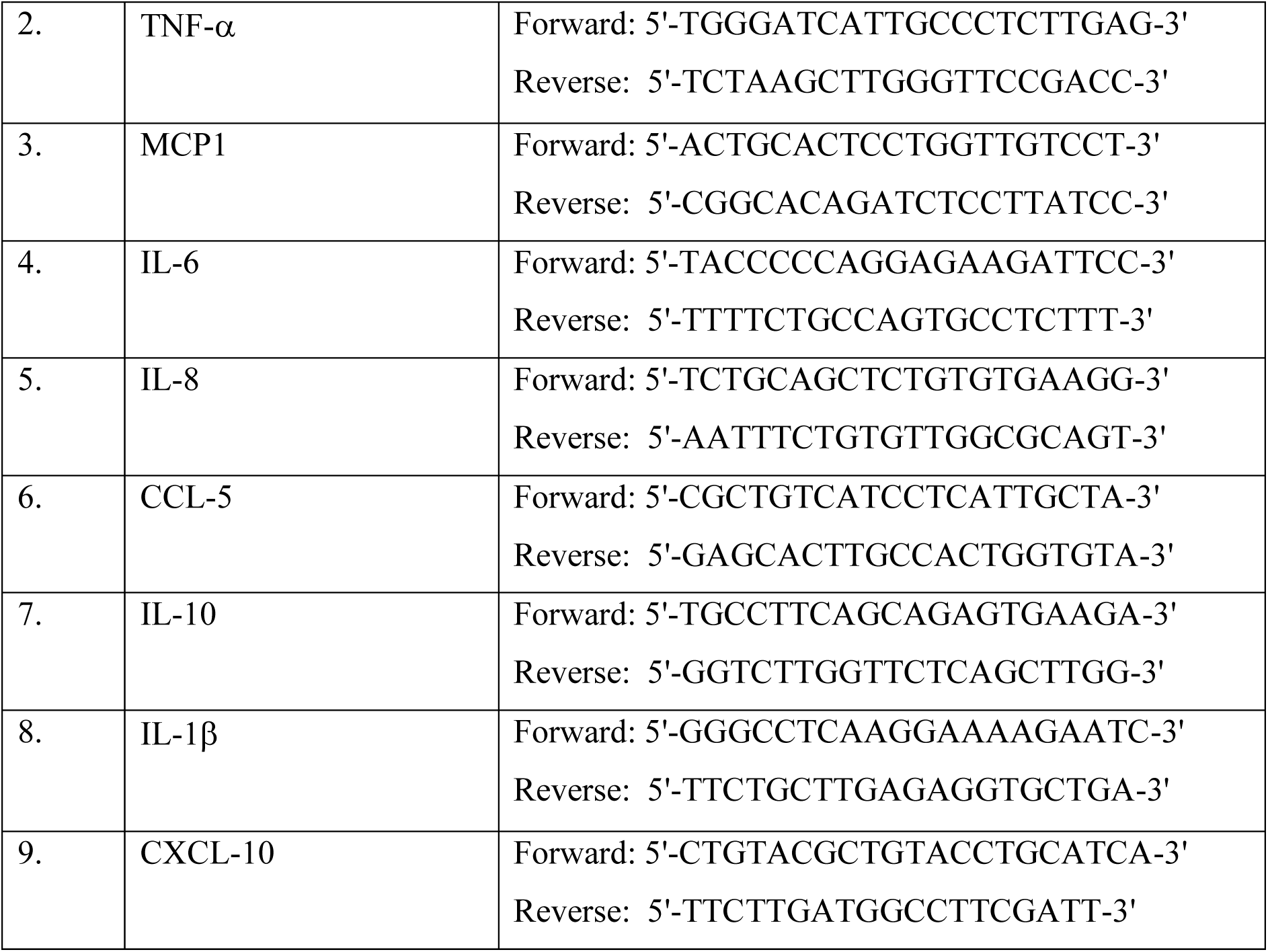
Human specific sequences of Forward and Reverse primers for various inflammatory genes.

### Cytokine measurement in the culture supernatants

Inflammatory cytokines/chemokines levels were determined in the culture supernatant using ELISA kits as per the manufacturer’s instructions (Invitrogen, Thermofisher Scientific). Sample dilutions were calibrated for respective analyte separately prior to analysing test samples. The reaction fluorescence was measured at 450 and 570nm wavelength (Molecular devices, Spectramax M2).

### Statistical Analysis

All experiments were carried out independently in triplicate. The statistical analysis was done by one-way ANOVA with the Tukey test for multiple comparisons and a two-tailed unpaired Student’s t-test for comparisons between two groups on GraphPad Prism (Version 9). Data are expressed as mean ± SEM. A p-value of less than 0.05 between the compared groups was considered statistically significant.

## RESULTS

Given the pressing need for immunosuppressants derived from plants, which offer favourable safety profiles and the ability to regulate the inflammatory response without compromising overall immune function, our study aims to evaluate the efficacy of three herbal extracts (WS, TC, GG) and an herbal formulation (A-64) in restoring the dysregulated inflammatory environment within a TLR7/8-induced macrophage cell model. As a reference compound, dexamethasone was included to assess the sensitivity of our model. In our experimental approach, macrophage-like cells were differentiated and pre-treated with aqueous extracts of the selected herbs at various concentrations (ranging from 3μg/mL to 300μg/mL) prior to TLR induction using either IMQ or RSQ. The induced cells exhibited a significant increase in the expression of inflammatory markers. Conversely, the cells treated with dexamethasone demonstrated improvement in the expression of all the investigated inflammatory genes. This investigation aims to shed light on the potential of these herbal extracts and formulation to modulate the dysregulated inflammatory response observed in our TLR7/8-induced macrophage cell model. By assessing their impact on the expression of inflammatory markers, we can gain insights into their efficacy as immunosuppressants. Notably, our study highlights the promising role of plant-derived compounds in restoring the balance of the inflammatory milieu without compromising the immune system’s ability to mount an appropriate response.

### Titration of agonist concentration and treatment duration

To determine the appropriate dose and induction duration for herbal evaluation, inflammatory genes (TNF-α, MCP1, IL-6, IL-1β) were examined in cells and culture supernatants treated with TLR agonists (IMQ and RSQ). Various concentrations (1µg/mL, 5µg/mL, and 10µg/mL) of agonists were tested in differentiated macrophage-like cells for 8, 16, 24, and 36 hours. IMQ induced TLR7/8 signaling after 8 hours, while RSQ showed induction after 24 hours. IMQ had maximum induction at 5µg/mL, whereas RSQ induced signaling at 1µg/mL. Thus, separate induction profiles were designed for herbals: 8 hours with 5µg/mL IMQ and 24 hours with 1µg/mL RSQ. TNF-α, MCP1, and IL-1β showed significant induction with 5µg/mL IMQ after 8 hours, while IL-6 showed induction after 16 hours (Supplementary Figure 2 and 3).

Before studying the effect of herbals on TLR signaling, their impact on cell viability was assessed using MTT assay, both in the absence and presence of TLR7/8 agonists. Even at a high concentration of 1000µg/mL, none of the herbs showed any harmful effects on cell viability (Supplementary Figure 1). Subsequently, the efficacy of selected herbal extracts in mitigating IMQ/RSQ-induced inflammatory responses in macrophage cells was investigated. The study design involved inducing TLR signaling with agonists, resulting in increased expression/circulating levels of inflammatory markers, and assessing the effect of the herbals on these markers.

### Prophylactic effect of *W. somnifera* (WS) on TLR7/8-induced Inflammatory markers

*W. somnifera*, commonly known as Ashwagandha, is an Indian shrub used in Ayurvedic medicine for holistic health benefits, stress and anxiety management, reproductive health support, physical endurance enhancement, and immunomodulation [2,35,14]. WS showed suppression of pro-inflammatory cytokine-induced Th1, Th2, and Th17 differentiation. It created an immunosuppressive environment in hamster and hACE2 transgenic mice models of COVID-19, suggesting potential efficacy against acute viral infections. WS had a more pronounced prophylactic effect in RSQ-induced cells compared to IMQ-induced cells (Figure 1). Increasing WS concentration significantly corrected the expression levels of inflammatory genes (Figure 1B). WS effectively mitigated TNF-α, CXCL10, and IL-1β expression in both IMQ and RSQ-induced macrophage-like cells (Figure 1A). Interestingly, WS treatment decreased IL-10 levels in RSQ-induced cells but had no such effect after IMQ induction. WS also significantly reduced MCP1 and IL-6 levels in RSQ-induced cells (Figure 1B). ELISA results showed significant reduction of MCP1 levels in culture supernatants of WS-treated IMQ-induced cells across all concentrations (Figure 2A). Treatment with 300µg/mL WS mitigated the effect of RSQ induction on IL6 and IL10 (Figure 2B). No correction in levels of TNF-α was observed in the supernatant of WS-treated cells induced with either agonist.

**Figure 1.**
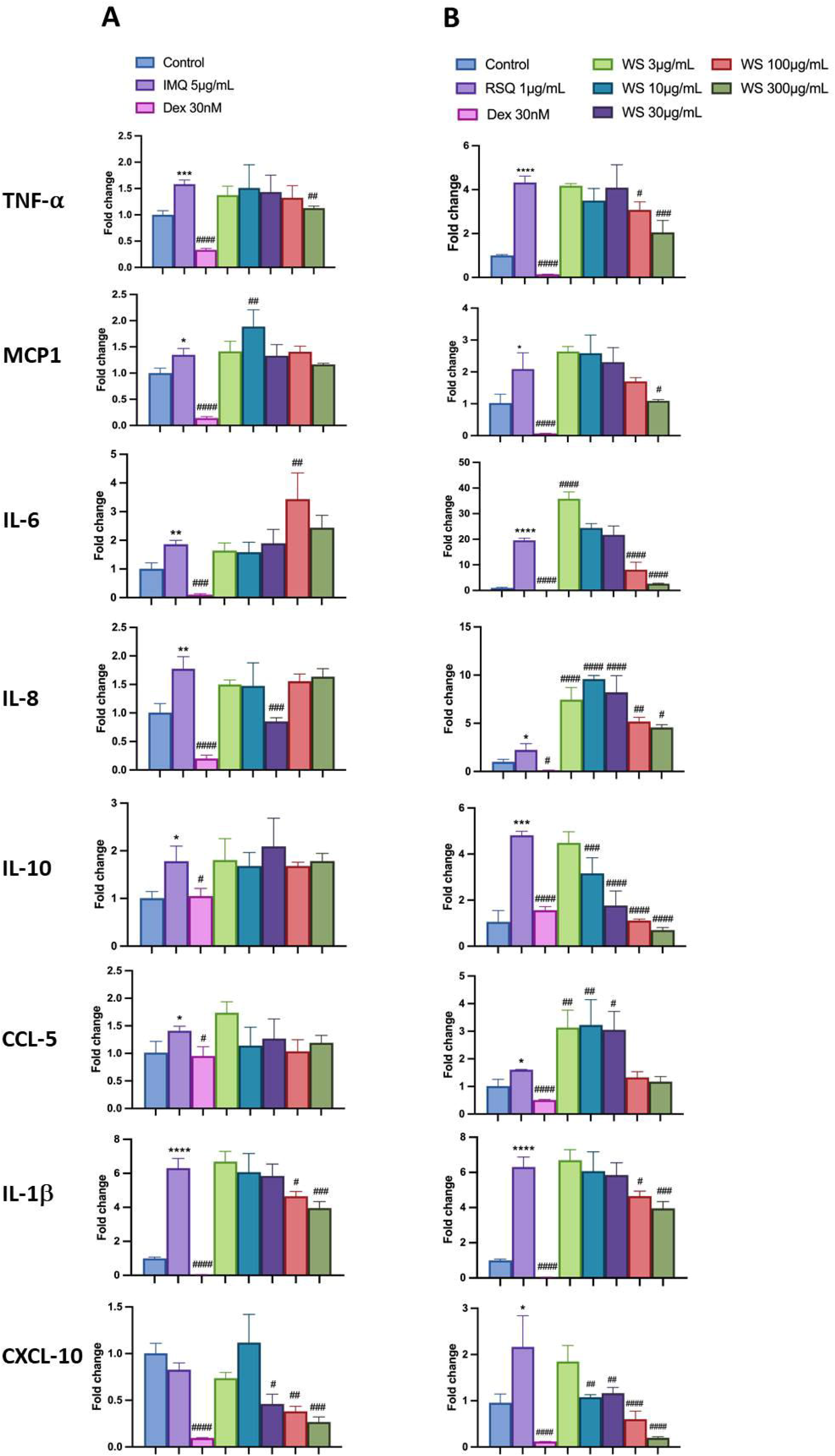
Concentration-dependent prophylactic effect of WS on the TLR7/8-induced gene expression of inflammatory markers. The cells were pretreated with WS for at least 30 minutes at 300µg/mL, 100µg/mL, 30µg/mL, 10µg/mL, and 3µg/mL concentrations and subsequently induced with IMQ (A)/ RSQ (B). Dexamethasone (30nM) was used as a reference control for the assay. All p-values were determined with respect to IMQ/RSQ-induced untreated group as well as the uninduced untreated group. All data are expressed as mean ± SEM.

**Figure 2.**
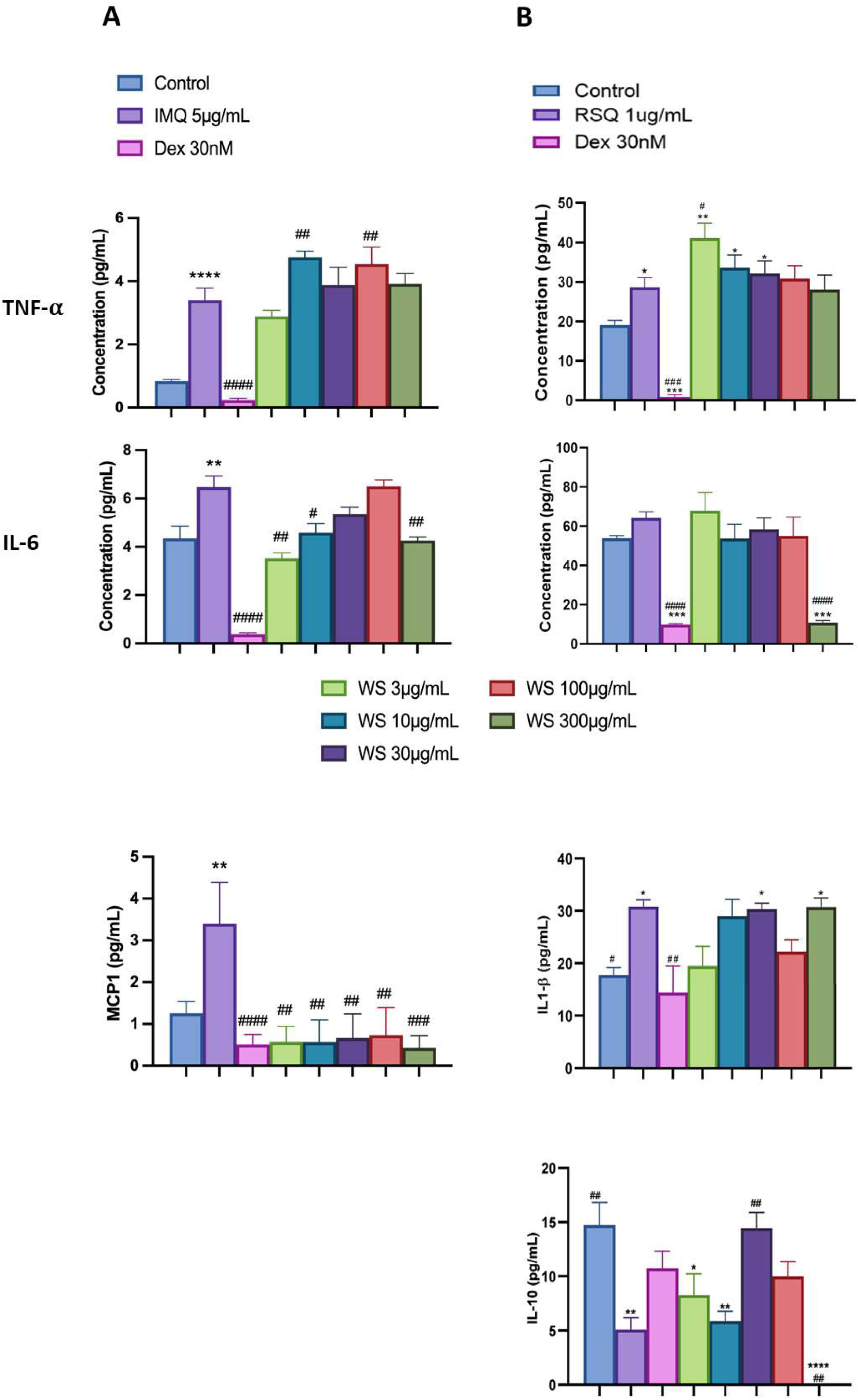
Concentration-dependent effect of WS on TLR7/8 induced secreted levels of cytokines/chemokines. The effect of WS treatment is portrayed on secreted levels of inflammatory markers in IMQ-induced (A) and RSQ-induced (B) macrophage-like cells. All p-values were calculated with respect to IMQ/RSQ-induced untreated group as well as the uninduced untreated group. Dexamethasone (30nM) was used as a reference control. Samples were diluted 50 times for TNF-α and MCP1 cytokines. All data are expressed as mean ± SEM

### TLR7/8-induced Inflammatory markers in *T. cordifolia* (TC) pretreated cells

*T. cordifolia* (TC), also known as Giloy or Guduchi, is a perennial shrub native to tropical regions of India and other neighbouring countries. In Ayurveda, the stem, leaves, and roots of TC have long been used for therapeutic purposes. It exhibits immunomodulatory properties, supporting the body’s defense against infection, and possesses antioxidant activity to reduce stress. Additionally, it offers anti-inflammatory, antipyretic, and hepatoprotective effects [29]. In-silico studies suggest that its active phytoconstituents bind to RNA-dependent RNA polymerase [34].

The prophylactic effect of TC on IMQ/RSQ-induced macrophage-like cells showed significant impact on inflammatory markers. After 24 hours of treatment with 300µg/mL TC, differentiated macrophages exhibited a noteworthy reduction in RSQ-induced expression of TNF-α, MCP1, IL-6, IL-10, CCL-5, and CXCL-10 (Figure 3B). Similar to WS, TC demonstrated a dose-response relationship, with increasing TC concentrations leading to significant reductions in IL-10 and CXCL-10 expression. Notably, ELISA results of TC-treated cell culture supernatants revealed interesting findings. While RSQ-induced IL-6 levels were mitigated at all investigated TC doses, the maximum decrease was observed at a minimal TC dose of 3µg/mL (Figure 4B).

**Figure 3.**
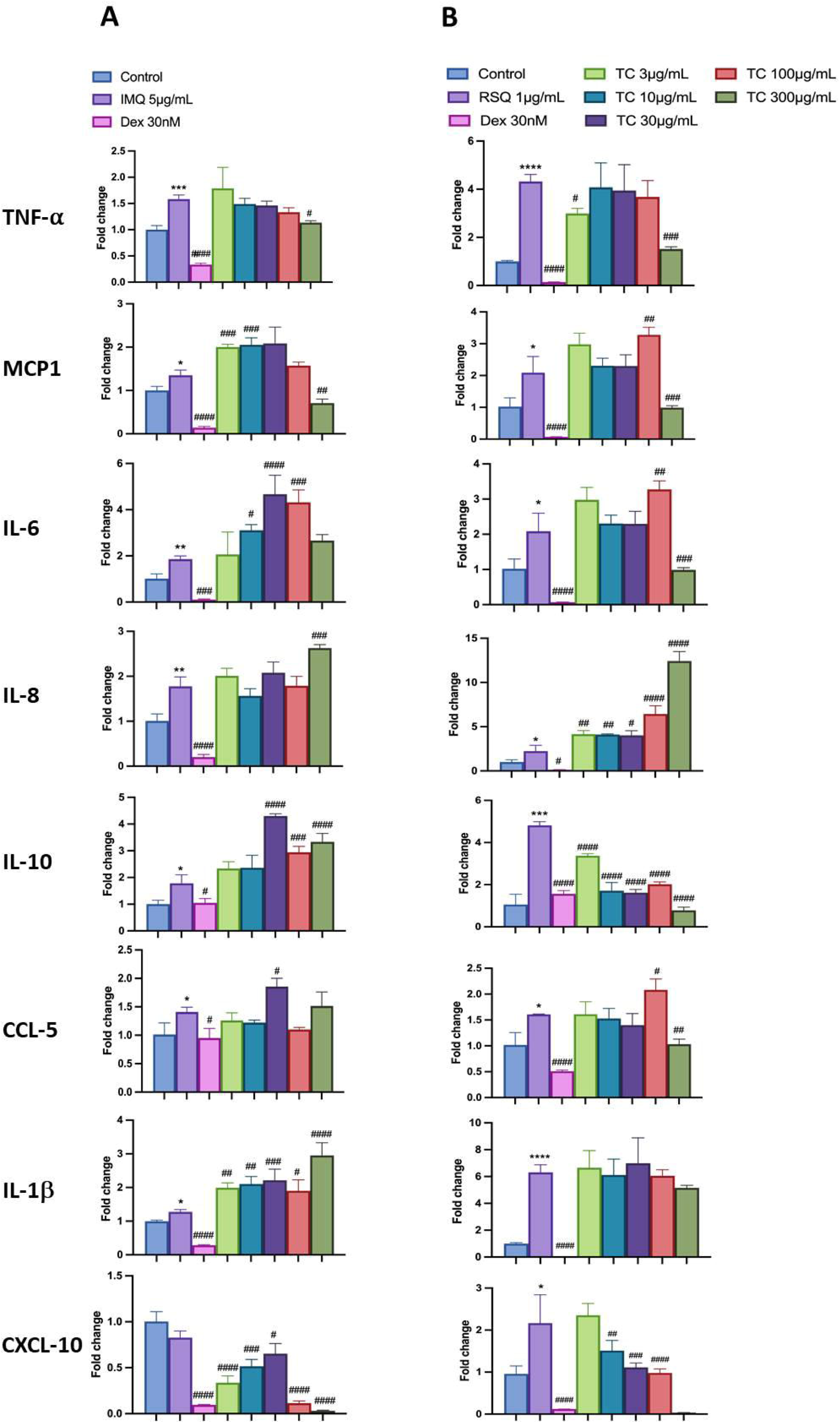
Concentration-dependent prophylactic effect of TC on the TLR7/8-induced gene expression of inflammatory markers. The cells were pre-treated for at least 30 minutes in the presence of TC at 300µg/mL, 100µg/mL, 30µg/mL, 10µg/mL, and 3µg/mL concentrations and subsequently induced with either IMQ (A) or RSQ (B). Dexamethasone (30nM) was used as a reference control for the assay. All p-values were determined relative to IMQ/ RSQ-induced untreated group as well as the uninduced untreated group. All data are expressed as mean ± SEM.

**Figure 4.**
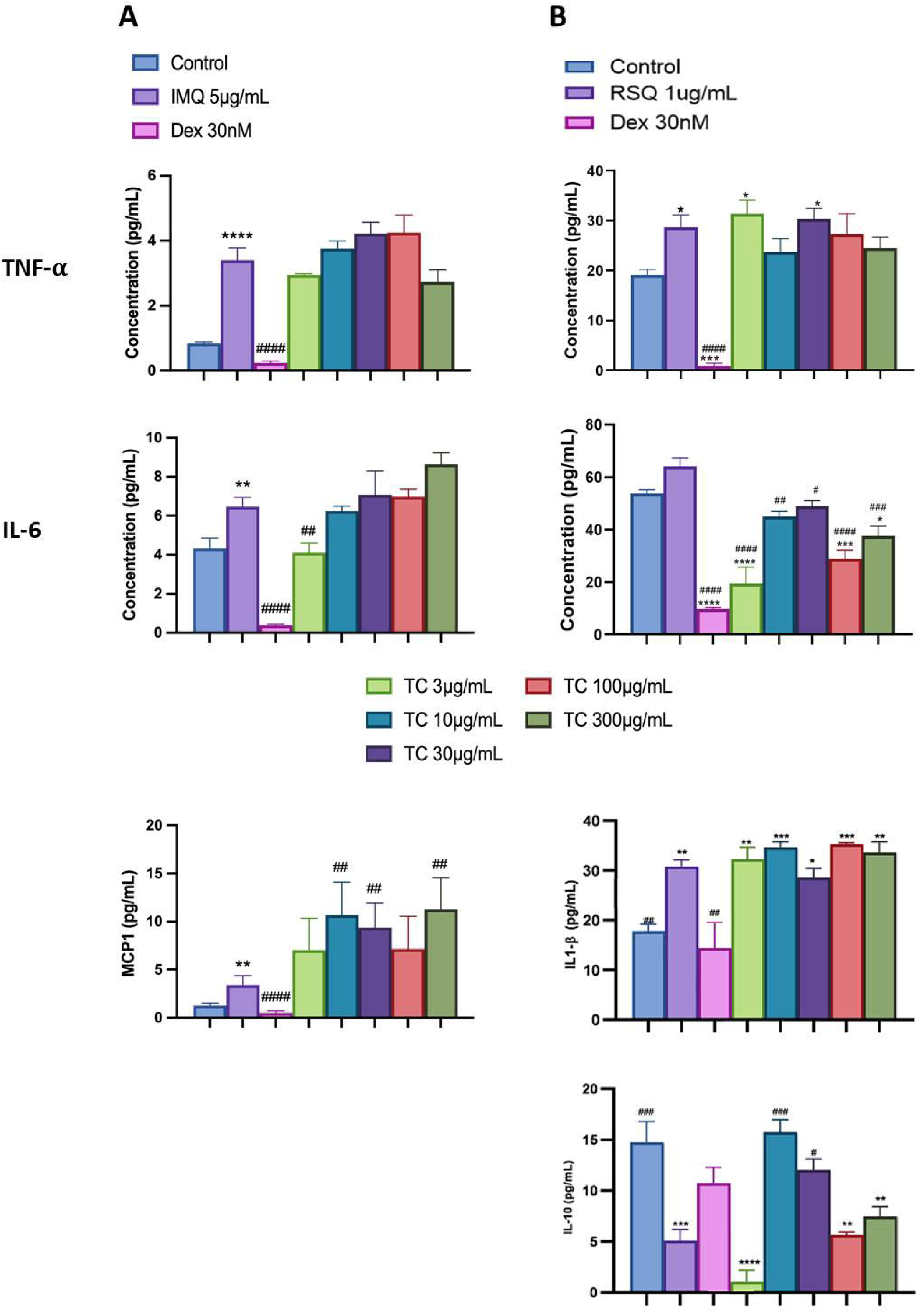
ELISA results of concentration-dependent prophylactic effect of TC on TLR7/8 induced secreted levels of inflammatory markers: The bar graph depicts the effect of TC treatment on secreted levels of different cytokines in IMQ-induced (A) and RSQ-induced (B) macrophage-like cells. All p-values were calculated with respect to IMQ/RSQ-induced untreated group as well as the uninduced untreated group. Dexamethasone was used as a reference control. Sample dilution was optimized to 50-fold to ensure optimal measurement of TNF-α and MCP1 cytokines. All data are expressed as mean ± SEM.

### Effect of *G. glabra* (GG) pretreatment on TLR7/8-induced Inflammatory markers

Popularly known as liquorice or mulaithi, GG is an ancient Ayurvedic herb widely used for its medicinal properties. GG roots contain active compounds like glycyrrhizin, flavonoids, coumarins, and triterpenoids, contributing to its anti-inflammatory effects in arthritis, expectorant properties for respiratory congestion, gastric and digestive benefits, antimicrobial properties, and immune regulation during infections [21]. GG administration in hamster models of COVID-19 demonstrated significant downregulation of pro-inflammatory cytokines TNF-α and IL-6 [27,37].

In our study, GG treatment led to a notable reduction in TNF-α and IL-6 expression and levels in macrophage cells induced with TLR7/8 agonists. IL6 gene expression significantly decreased at 3µg/mL GG concentration but increased with higher GG concentrations. IL-8 gene expression was inhibited at 300µg/mL, 30µg/mL, and 10µg/mL, while CXCL10 gene expression decreased in a concentration-dependent manner in IMQ-induced cells, with the maximum reduction observed at 300µg/mL GG (Figure 5A). GG treatment significantly reduced TNF-α expression at all investigated concentrations. Similarly, GG treatment corrected IL-10 expression in RSQ-induced cells (Figure 5B). IL8 depletion was observed at 10µg/mL GG, while CCL5 gene expression was significantly corrected at lower GG concentrations (3µg/mL and 10µg/mL) in RSQ-induced cells. Surprisingly, IL1-β gene expression increased with increasing GG concentration in IMQ-induced cells. MCP1 gene expression showed an increasing trend in GG-treated cells induced with both TLR7/8 agonists. Likewise, circulating MCP1 protein levels increased in culture supernatants with higher GG concentrations. However, IL6 levels were significantly reversed at most GG concentrations investigated in RSQ-induced cells (Figure 6B).

**Figure 5.**
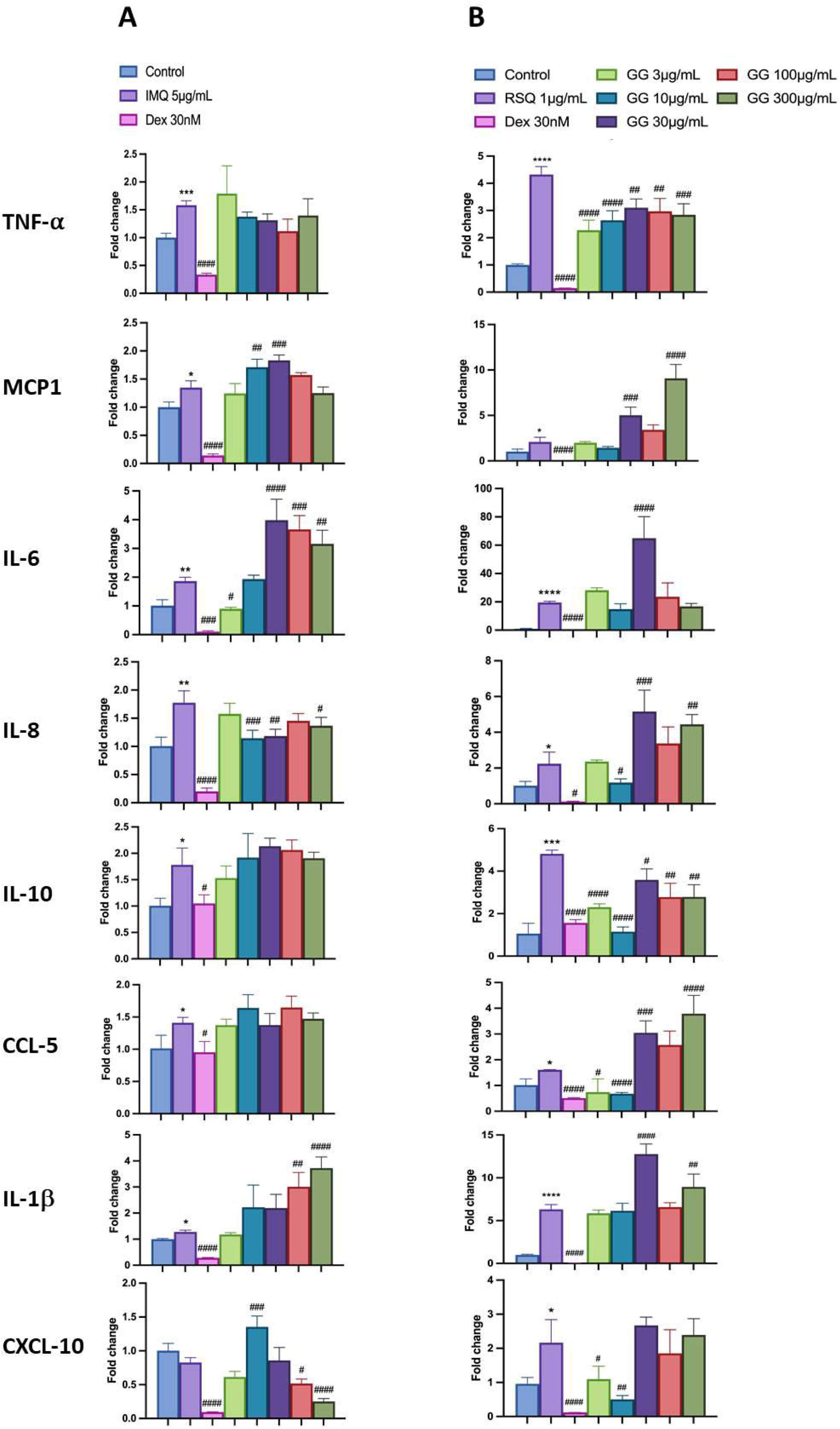
Concentration-dependent prophylactic effect of GG treatment on the TLR7/8 induced gene expression of inflammatory markers: The cells were pretreated with GG at 300µg/mL, 100µg/mL, 30µg/mL, 10µg/mL, and 3µg/mL concentrations and thereafter induced with IMQ (A)/ RSQ (B). Dexamethasone (30nM) was used as a reference control. All p-values were determined with respect to IMQ/ RSQ-induced untreated group as well as the uninduced untreated group. All data are expressed as mean ± SEM.

**Figure 6.**
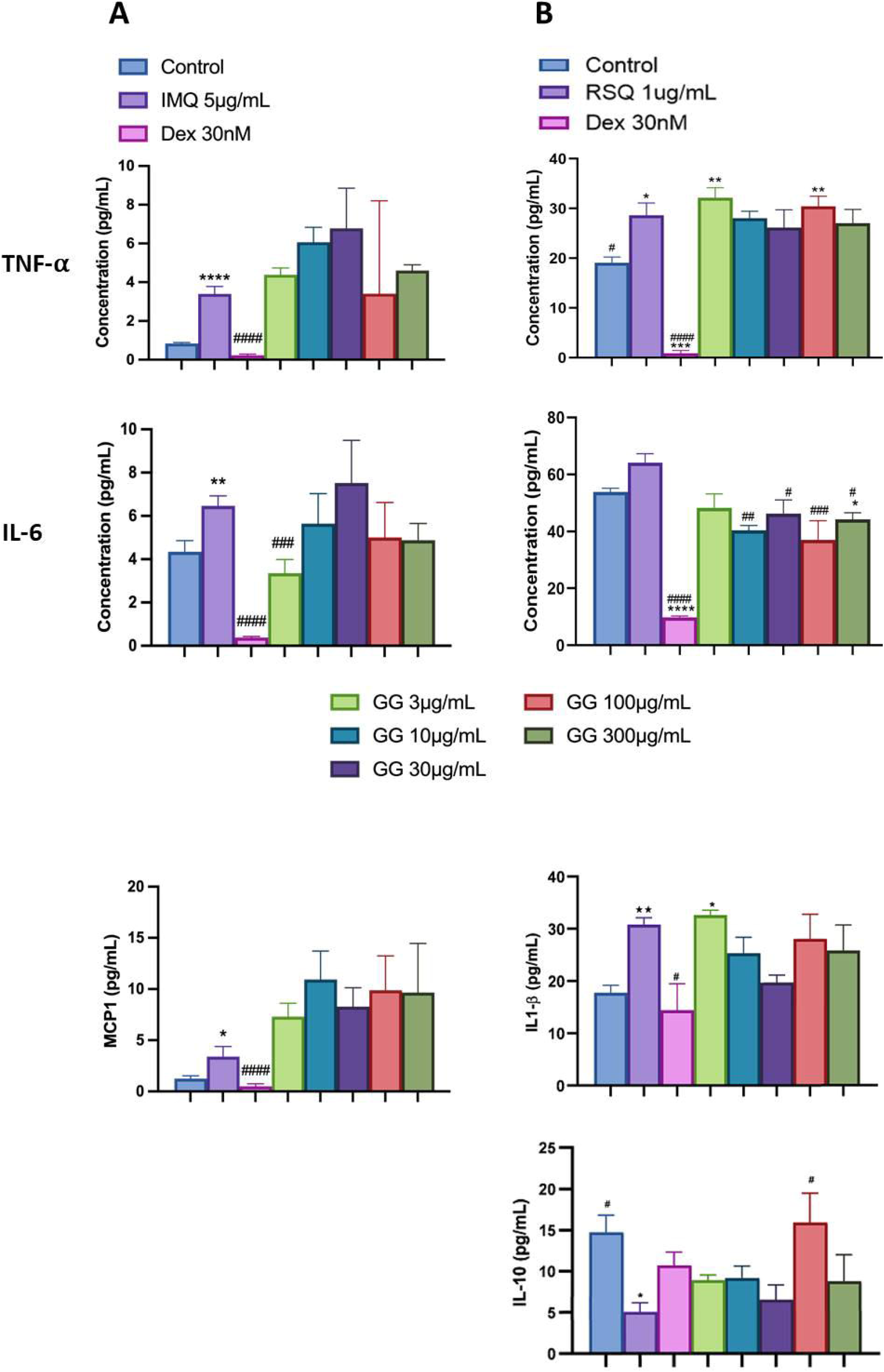
Concentration-dependent prophylactic effect of GG treatment on TLR7/8 induced secreted levels of inflammatory markers: The bar graph presents changes in secreted levels of inflammatory markers after treatment with a range of GG concentrations in IMQ-induced (A) and RSQ-induced (B) macrophage-like cells. All p-values were calculated with respect to IMQ/RSQ-induced untreated group as well as the uninduced untreated group. Dexamethasone was used as a reference control. Sample dilution was calibrated to 50-fold for optimal measurement of TNF-α and MCP1 cytokines. All data are expressed as mean ± SEM.

### Effect of the pretreatment of cells with AYUSH-64 (A-64) on the TLR7/8 induced expression and release of inflammatory markers

AYUSH-64 is an Ayurvedic formulation with immunomodulatory and anti-inflammatory properties. It contains herbs such as *Alstonia scholaris*, *Caesalpinia crista*, *Picrorhiza kurroa*, and *Swertia chirata* [23]. Safety studies have confirmed its tolerance up to 500mg/kg body weight in rodents. A clinical trial conducted by the Ministry of AYUSH, India, and the Council of Scientific and Industrial Research, India, demonstrated its efficacy in managing mild to moderate COVID-19 patients, with a significant decrease in IL-6 levels [31,32,25]. In our study, we evaluated the effect of A-64 on the expression of inflammatory genes in the presence of TLR7/8 inducers (Figure 7). We observed a significant reduction in TNF-α expression and levels in the supernatant of A-64 pre-treated cells. While, IL6 levels showed no overall change in expression or in the release in the supernatant, IL-1β levels decreased significantly with increasing A-64 concentration, indicating a concentration dependent-response relationship (Figure 8). Moreover, IL-10 and CXCL10 expression were mitigated in a concentration-dependent manner, decreasing with increasing AYUSH-64 concentrations.

**Figure 7.**
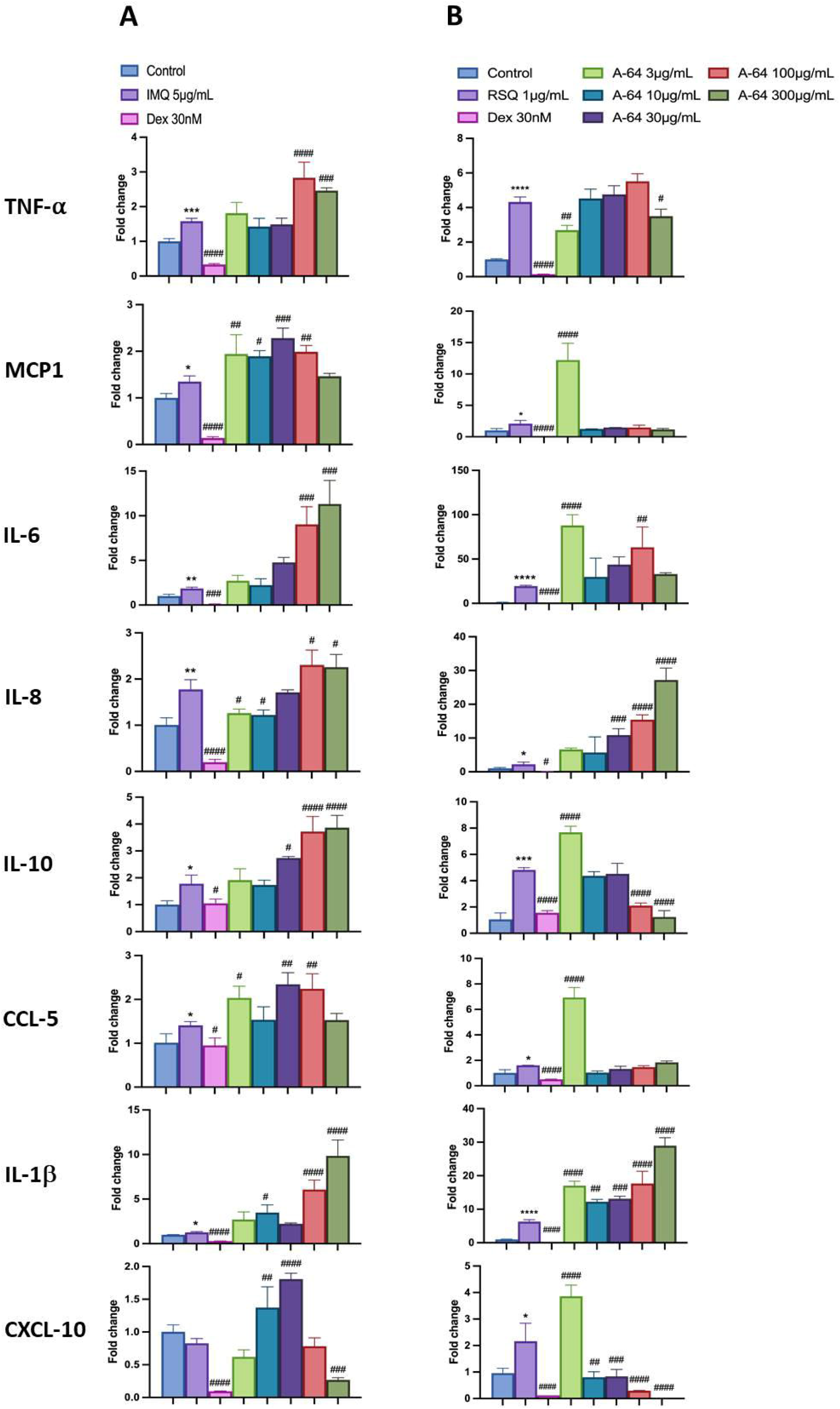
Concentration-dependent prophylactic effect of A-64 treatment on the TLR7/8 induced gene expression of inflammatory markers: The cells were pretreated for at least 30 minutes with A-64 at 300µg/mL, 100µg/mL, 30µg/mL, 10µg/mL, and 3µg/mL concentrations and later induced with IMQ (A)/ RSQ (B). Dexamethasone (30nM) was used as a reference control for the alleviation of TLR7/8 response. All p-values were determined with respect to IMQ/ RSQ-induced untreated group as well as the uninduced untreated group. All data are expressed as mean ± SEM.

**Figure 8.**
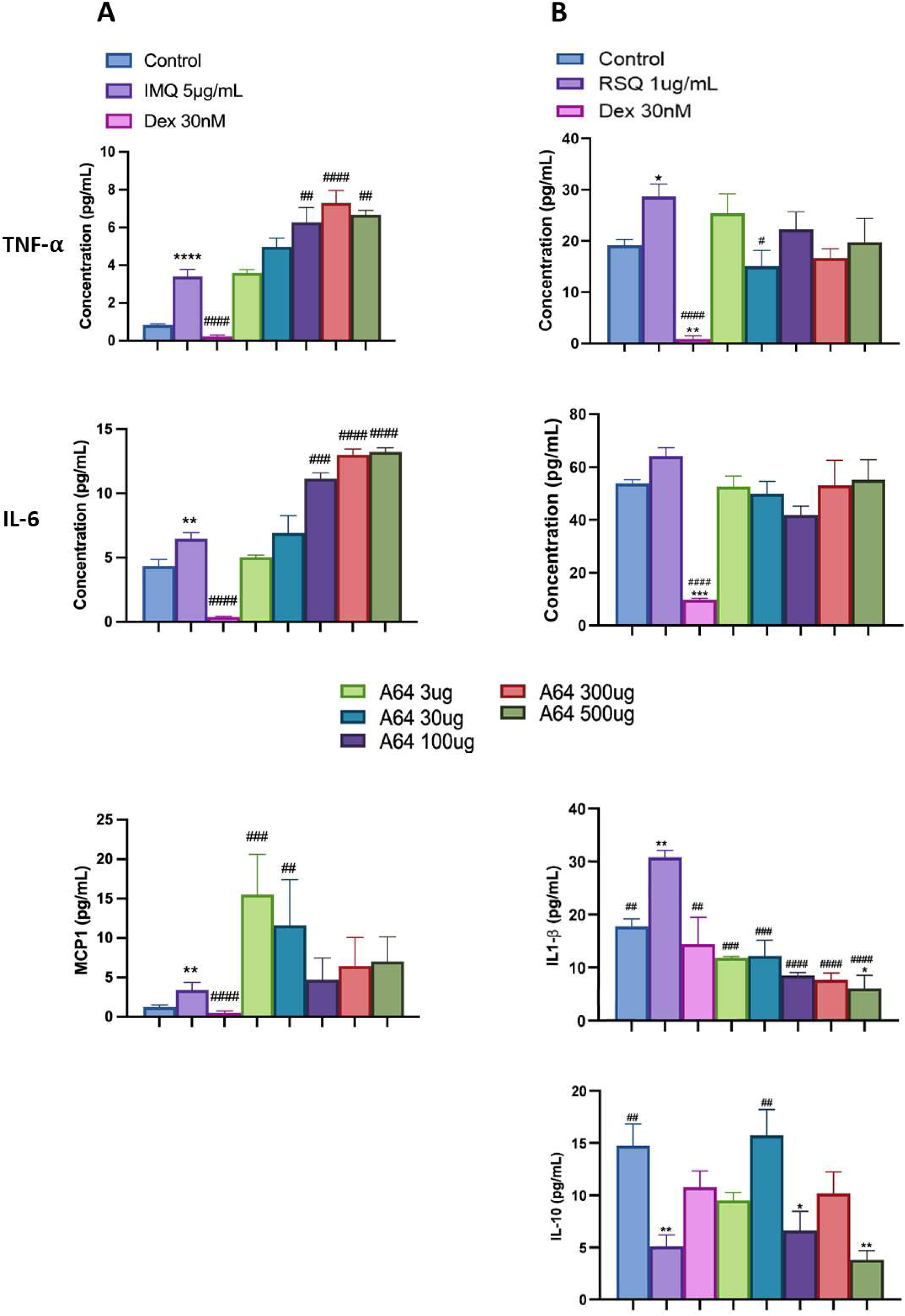
Prophylactic effect of A-64 treatment on TLR7/8 induced secreted levels on inflammatory markers: The bar graph depicts changes in secreted levels of different inflammatory markers as a result of treatment with AYUSH-64 herbal formulation in IMQ-induced (A) and RSQ-induced (B) macrophage-like cells. All p-values were calculated with respect to IMQ/RSQ-induced untreated group as well as the uninduced untreated group. Dexamethasone (30nM) was used as a reference control. Sample dilution was calibrated to 50-fold for the optimal determination of both TNF-α and MCP1 cytokines. All data are expressed as mean ± SEM.

Overall, our findings contribute to the growing body of research on plant-based therapeutics, particularly in the context of immunomodulation. The use of herbal extracts and formulations holds great potential in the development of effective treatments for inflammatory disorders, offering a safe and well-tolerated alternative to conventional immunosuppressive agents.

## DISCUSSION

The dysregulated inflammatory response has emerged as a prominent pathology during the recent COVID-19 pandemic. To gain a comprehensive understanding of the disease, cell-based models have played a crucial role in elucidating the mechanisms behind various candidate molecules in COVID-19 therapeutics. Specifically, the activation of Toll-like receptor 7/8 (TLR7/8) signaling pathway in response to the presence of viral RNA within host cells has been implicated in the release of pro-inflammatory cytokines, essential for mediating the anti-viral immune response. In our study, we employed macrophage-like differentiated THP1 cells as a model system to mimic TLR7/8 induction. This was achieved by treating the cells with imidazoquinolinamines, namely imiquimod (IMQ) and resiquimod (RSQ). IMQ and RSQ are well-known TLR7/8 agonists that have demonstrated potent ability to induce the production of interferon-α (IFN-α) and other cytokines [9,8].

By utilizing this experimental setup, we aimed to gain mechanistic insights into the TLR7/8-mediated inflammatory response and its potential implications in immune-compromised pathologies including COVID-19. The use of cell-based models has proven to be invaluable in unravelling the complex interplay between viral components and the host immune system, paving the way for the development of novel therapeutic strategies. By elucidating the specific mechanisms underlying herbal modulation of TLR7/8 signaling and subsequent cytokine release, our study contributes to a better understanding of the dysregulated inflammatory response observed during COVID-19. These findings may have implications for the development of targeted therapeutics aimed at modulating the TLR7/8 pathway to mitigate the excessive inflammation associated with the disease.

Excessive and dysregulated release of pro-inflammatory cytokines, known as cytokine storm, can occur during severe infections and autoimmune disorders [10]. This uncontrolled cytokine release was observed in the recent COVID-19 pandemic, leading to life-threatening systemic inflammation. Immunomodulatory therapies targeting specific cytokine pathways, such as TNF-α, IL-6, and IL-1β, are used to manage cytokine storm [18]. Tocilizumab, an IL-6 receptor inhibitor, has shown efficacy in this regard [38]. IL-1β, on the other hand, induces IL-6 and TNF-α production and activates immune cells [5]. Elevated levels of TNF-α, IL-6, and IL-1β are associated with severe symptoms and tissue damage. Blocking IL-1 signalling with antagonists like anakinra has potential in managing the inflammatory response [15]. However, synthetic immunomodulatory agents often come with adverse side effects [6]. In contrast, the herbs investigated in this study (WS, TC, GG, A-64) have a long history of traditional use. Their anti-inflammatory and immunomodulatory properties offer a potential approach to manage the hyperinflammatory response without toxic side effects [11,4]. These botanical sources of immunomodulatory agents provide a novel avenue to moderate the exaggerated cytokine response and restore immune balance [11,4].

TLR7/8 activation plays a crucial role in recruiting immune cells to the site of viral infection by triggering pro-inflammatory cytokine production [19,9,1]. However, the administration of TLR7/8 agonists like imiquimod or resiquimod can lead to cytokine storms in certain clinical scenarios. Macrophage-like THP1 cells, which differentiate from monocytes when exposed to PMA, provide a relevant model to study the dysregulated cytokine response induced by TLR7/8 agonists and evaluate the prophylactic effect of herbal extracts. The study model was optimized for agonist dosage and treatment duration to achieve optimal TLR signaling induction. RSQ showed a higher induction rate of cytokine expression compared to IMQ, and the concentration of RSQ required for TLR7/8 induction (1µg/mL) was five times lower than that of IMQ (5µg/mL). Similar observations were reported by another research group using a similar model [8].

Induction of TLR7/8 signaling with agonists increased TNF-α expression, which was moderated in presence of dexamethasone. Treatment with herbal extracts significantly improved TNF-α gene expression in macrophage-like cells, with expression generally decreasing as extract concentration increased. Among the extracts, WS and TC showed significant correction of IL-6 gene expression, while all treatment groups exhibited perturbed levels of IL-6 in the culture supernatant. WS treatment also corrected TLR7/8-induced IL-1β levels, whereas other extracts had no effect on IL-1β gene levels. The interplay between IL-1β and IL-6 regulation is crucial for understanding inflammatory processes and developing therapeutic strategies [3]. A-64 inhibition of IL-1β resulted in suppressed IL-6 release. Additionally, the selected herbs significantly corrected CXCL-10 expression, an immune cell trafficking and activation chemokine. CXCL-10 levels are associated with disease severity and can serve as a biomarker in inflammatory conditions [16].

Among all the investigated herbal extracts, W. somnifera (WS) showed remarkable efficacy in mitigating the increase in expression of inflammatory markers induced by TLR7/8 activation. WS exhibited its effects by attenuating excessive cytokine release and regulating the production and activity of pro-inflammatory cytokines. WS contains bioactive compounds, including withanolides, known for their anti-inflammatory and immunomodulatory properties. Previous studies have demonstrated the ability of WS to suppress TNF-α, IL-6, and IL-1β cytokines in various inflammatory conditions, such as LPS-induced neuroinflammation [12]. Furthermore, WS has shown a protective effect against COVID-19 by modulating the inflammatory response and inhibiting pro-inflammatory cytokine-induced differentiation of Th1, Th2 and Th17 cells [28].

It is essential to acknowledge that although these botanical drugs have demonstrated promising results in our preclinical model, further research is warranted to comprehensively understand their mechanisms of action. Additionally, optimizing dosages and conducting thorough evaluations of their safety and efficacy in animal models are crucial steps to advance their potential as therapeutic interventions. While our study model exhibits sensitivity and robustness, rigorous clinical validation is imperative to establish the viability and effectiveness of these botanical drugs as potential treatment options.

## CONCLUSIONS

Based on our study, it is evident that herbal therapeutics hold promise in regulating dysregulated inflammatory responses. Resiquimod exhibited better efficacy as an agonist for activating the TLR7/8 signaling pathway in our experimental model. Among the evaluated herbs/formulations, *Withania somnifera* demonstrated significant prophylactic efficacy by effectively mitigating the upregulation of numerous inflammatory genes in the PMA-differentiated THP1 cells activated via the TLR7/8 signaling pathway. These findings provide a basis for exploring the potential of herbal interventions in modulating inflammatory cascades, but additional research is required to fully understand their mechanisms of action and therapeutic applications.

## FIGURE LEGENDS

**Supplementary Figure 1**

**Effect of AYUSH extracts on cell viability:** Bar graph depicting the effect of the investigated herbal extracts at 1000µg/mL concentration. Dexamethasone (1µM) was included as reference control as it was used in subsequent efficacy experiments at significantly lower concentrations to mitigate excess cytokine response in IMQ/RSQ induced macrophage-like cells. The data are expressed as mean ± SEM.

**Supplementary Figure 2**

**Time-dependent effect of TLR7/8-induced cytokine gene expression**: Inflammatory cytokines including TNF-α, MCP1, IL-6, IL-1β, and CXCL10 genes were investigated for changes in their gene expression after induction with IMQ (A) and RSQ (B) at three different concentrations (1µg/mL, 5µg/mL and 10µg/mL) over a range of time (8, 16, 24 and 36 hours). All p-values were determined with respect to untreated control cells. All data are expressed as mean ± SEM.

**Supplementary Figure 3**

**Temporal effect of TLR7/8-induced secreted cytokine levels**: Levels of TNF-α, MCP1, IL-6, and IL-1β cytokines investigated in 5µg/mL IMQ (A) and 1µg/mL RSQ (B) induced macrophage-like cells at three different time points – 8, 16 and 24 hours. All p-values were calculated with respect to untreated control cells. All data are expressed as mean ± SEM.

## Supporting information

Effect of AYUSH extracts on cell viability

Time-dependent effect of TLR7/8-induced cytokine gene expression

Temporal effect of TLR7/8-induced secreted cytokine levels

## Acknowledgments

The authors express their gratitude to the Ministry of AYUSH and the Department of Biotechnology (DBT), Government of India for jointly funding the research work presented in this manuscript (Grant Nos.: BT/PR40738/TRM/120/486/2020 and A.11019/03/2020-NMPB-IV-A). Madhu Dikshit also acknowledges the financial support from JBR/2020/000034. We acknowledge THSTI Small Animal Facility (SAF) for its services. The authors are thankful to NMPB for providing Herbal extracts and Formulations for the study. The authors thank the Executive Director of THSTI for his generous overall support. We also appreciate the technical assistance provided by Ms. Deepika Tuli from THSTI during *in-vitro* experimental procedures.

## Author contributions

**Manisha Dagar and Kamala Priya** performed the *in vitro* cell-based experiments, writing, and interpretation. **Ajay Kumar** and **Madhu Dikshit** conceptualized the overall study design, interpretations, writing, and editing of the manuscript. All authors have read and accepted the manuscript.

