## Supplementary figures and images for "Pharmacological Evaluations of Select Herbal Extracts on TLR7/8-induced Cytokine and Chemokine Production in Macrophage-like Cells"

### Effect of AYUSH extracts on cell viability

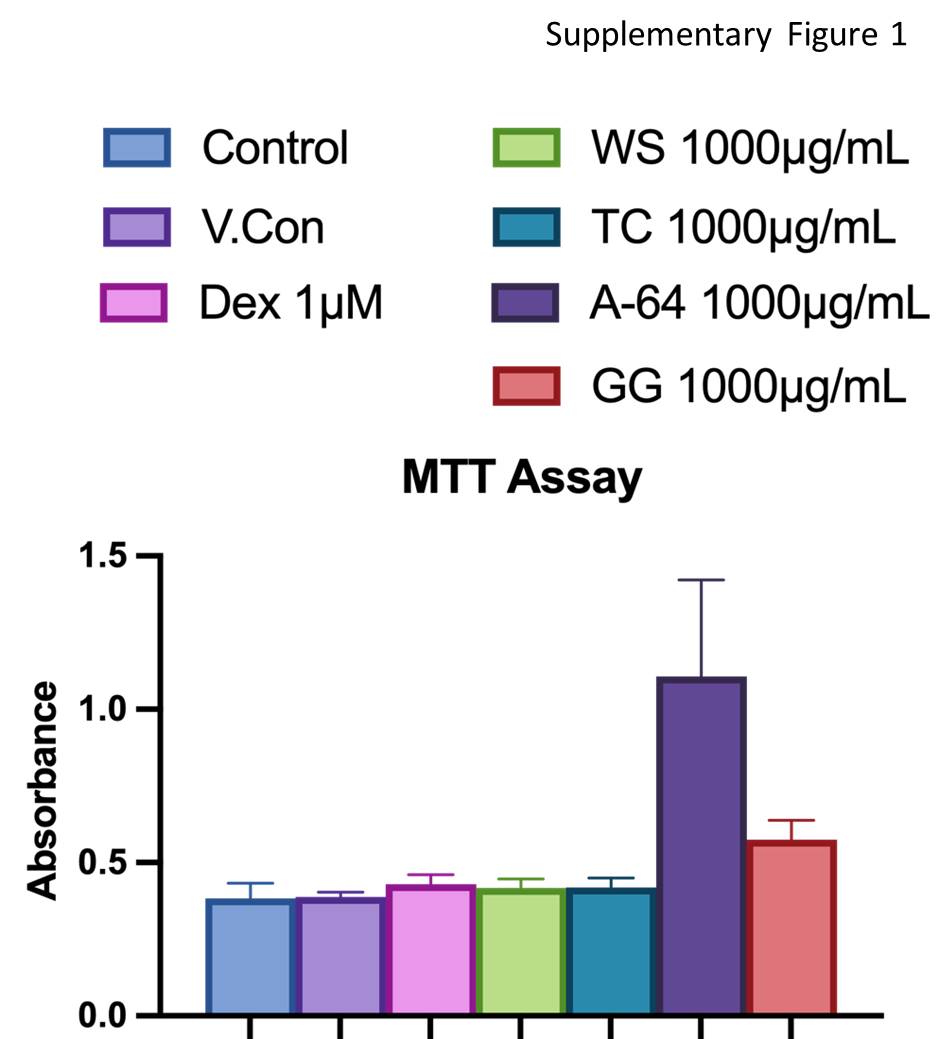

### Temporal effect of TLR7/8-induced secreted cytokine levels

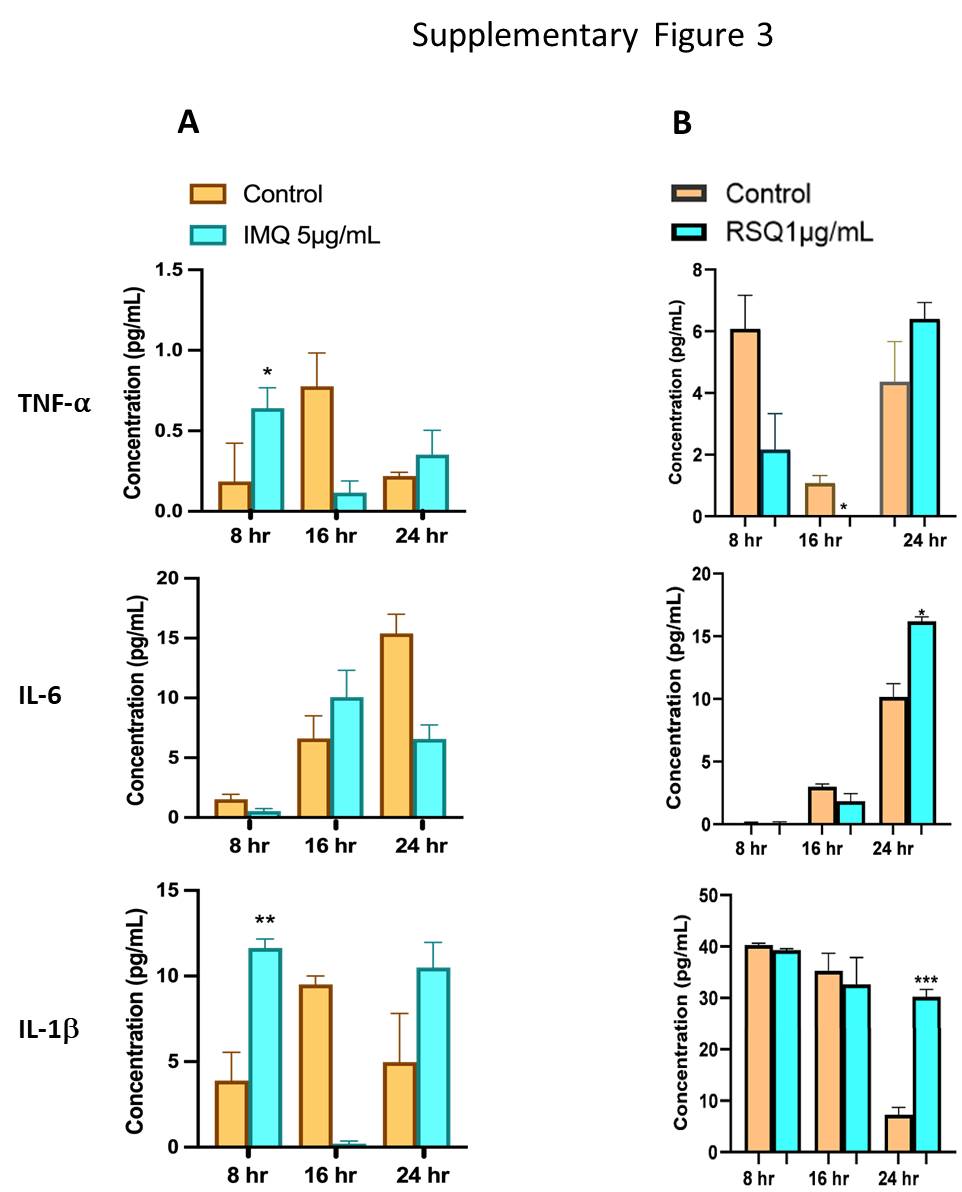

### Time-dependent effect of TLR7/8-induced cytokine gene expression

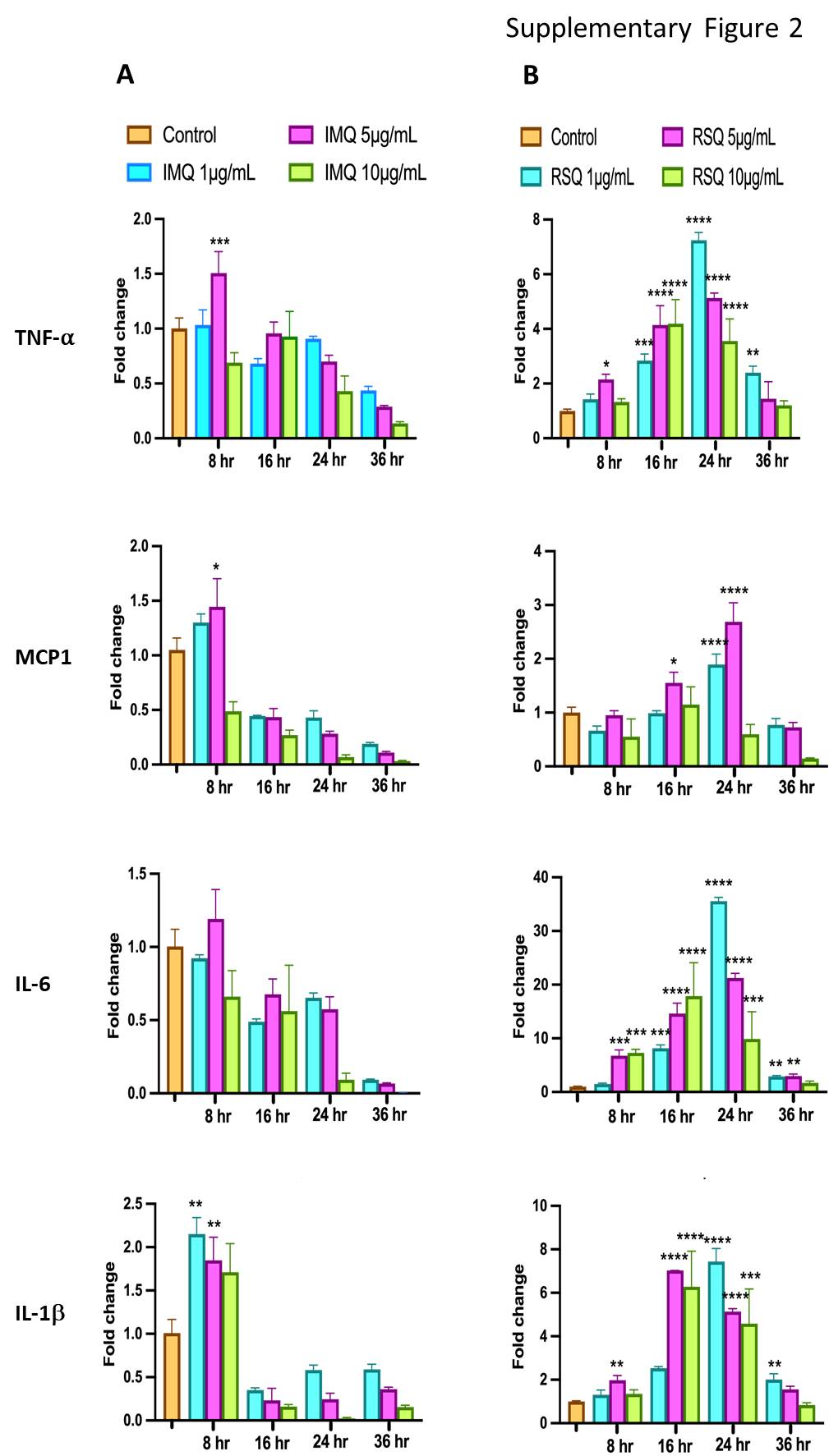
